# A highly efficient CRISPR-Cas9-based gene editing system in oat (*Avena sativa*)

**DOI:** 10.1101/2025.01.26.633040

**Authors:** Mehtab-Singh, Cali Kaye, Rajvinder Kaur, Jaswinder Singh

**Affiliations:** Plant Science Department, 21111 Rue Lakeshore, McGill University, Montreal, QC H9X 3V9, Canada

**Keywords:** CRISPR/Cas9, Oat, Gene-editing, VRN, TLP, PEBP, Plant Transformation, Beta-glucan, CAPS Assay, Gene Knockout

## Abstract

Cultivated oat (*Avena sativa*) is an emerging cereal for healthy lives owing to its unique characteristics, such as high β-glucan and oil content, distinctive fatty acid composition, and gluten-free nature. The recent unravelling of the 12.5 Gb hexaploid oat genome underlined breeding barriers caused by ancestral translocations and inversions, leading to recombination suppression and pseudo-linkage further hindering conventional trait introgression. Over the past decade, the Clustered Regularly Interspaced Short Palindromic Repeats (CRISPR)-Cas9 system has been extensively used for crop improvement and functional genomics in all other cereals except oats. Its large repetitive genome with three sub-genomes, lack of efficient transformation, recalcitrant nature, and complex molecular screening due to gene redundancy have been major obstacles to gene editing success in oat. We report the first successful CRISPR-Cas9-based gene editing in oat in three genes — *AsTLP8, AsVRN3* and *AsVRN3D* with gene-editing efficiency of up to 41.1%. The gene-edited plants for all the genes carried deletions and/or one base insertion. Further analysis of *VRN3* T_1_ and T_2_ mutants revealed bent leaves in heterozygous knockouts (AACCdD), while an extended vegetative growth phase was seen in the T_1_ homozygous and biallelic mutants (aaccdd), accentuating the important role of *VRN3* in oat development. We are confident that this highly efficient oat gene editing system will pave the way for a deeper molecular understanding of this healthy cereal, deciphering oat’s functional genomics, and creating genetic diversity at the cold spots of recombination in oat.

---

Cultivated oat (*Avena sativa*), a cereal of worldwide importance, belongs to the *Aveneae* family of cereals, distinguishing it from wheat, rye, barley, and rice. This distinction explains its unique characteristics, such as high β-glucan and oil content, distinctive fatty acid composition, and gluten-free nature, making it suitable for both human consumption and animal feed. Since the 1870s, oat breeding programs have achieved remarkable improvements in various traits, including disease resistance, yield, milling quality, and β-glucan content. However, the process remains lengthy, labour-intensive, and cumbersome. The recent unravelling of the massive 12.5 Gb oat genome revealed breeding barriers owing to the ancestral large-scale translocations and inversions (Kamal et al., 2022). Hence, the traditional introgression of traits through breeding cannot be attained due to recombination suppression and pseudo-linkage at such genetic loci. This highlights the urgent need to employ the latest genome editing technologies for oat improvement, which holds the utmost potential for altering specific gene functions and introducing allelic diversity. Over the past decade, the Clustered Regularly Interspaced Short Palindromic Repeats (CRISPR)- Cas9 system has been extensively used for crop improvement and functional genomics in all other cereals except oats (Ahmar et al., 2024). Its large repetitive genome with three sub-genomes, lack of efficient transformation, recalcitrant nature, and complex molecular screening due to gene redundancy are major obstacles to gene editing success. We report the first successful CRISPR-Cas9-based gene editing in oat, generating deletions and insertions.

To test whether the CRISPR-Cas9 system can produce targeted gene editing in oat, different single guide RNAs (sgRNAs) were designed to target the *Thaumatin-like protein 8 (TLP8), Vernalization 3 (VRN3)*, and *Vernalization 3-D (VRN3D)* in hexaploid oat (Figure 1a, b and Figure S5). *TLP8* has been associated with β-glucan the primary heart-healthy fibre that makes oats valuable for human consumption (Singh et al., 2017). Concomitantly, *VRN3* is a member of the PEBP (Phosphatidylethanolamine-binding protein) family and has been mapped to a QTL associated with plant height, oil content and other crucial yield-related traits in oat. Intriguingly, *VRN3D* is in a recombination-suppressed region on the inverted 7D chromosome, making it an interesting candidate for gene editing (Tinker et al., 2022). Mature seeds of the hexaploid spring oat variety *Park* were selected for transformation due to the recent success in our lab with the introduction of *Ac/Ds* elements and the refinement of fatty acid composition using particle gun bombardment (Mahmoud et al., 2022; Zhou et al., 2024). Seeds were sterilized with bleach, and calli were produced for plant transformation. The transformed calli underwent three rounds of hygromycin selection (20mg/L) followed by regeneration and rooting of transgenic calli on the respective media supplemented with 5mg/L hygromycin (Figure 1h; Methods S1). Three individual constructs (pJDTLP8, pJDVRN3, and pJDVRN3D) were designed by cloning the gene-specific guides driven by wheat U6 promoter in the JD633 backbone with ubiquitin promoters for GRF-GIF chimera, *Cas9*, and *hygromycin (hpt)* (Debernardi et al., 2020) (Figure 1a,b and Figure S5). For the *TLP8* gene, a total of 100 calli were bombarded with the pJDTLP8 construct, producing 21 transgenic plants with a transformation efficiency of 21% (Figure S1). The guide flanking region of the *AsTLP8* was then amplified using *TLP8* gene primers (Table S1) and subjected to next-generation sequencing (NGS). The analysis confirmed seven T_0_ gene-edited plants reporting a highly efficient gene editing frequency of 41.1% (Figure 1f). Notably, the PCK-7 plant had a 4bp deletion in the *TLP8C*, an 11bp deletion and a 1bp (T) insertion in the *TLP8D* (Figure 1c).

**Figure 1.**
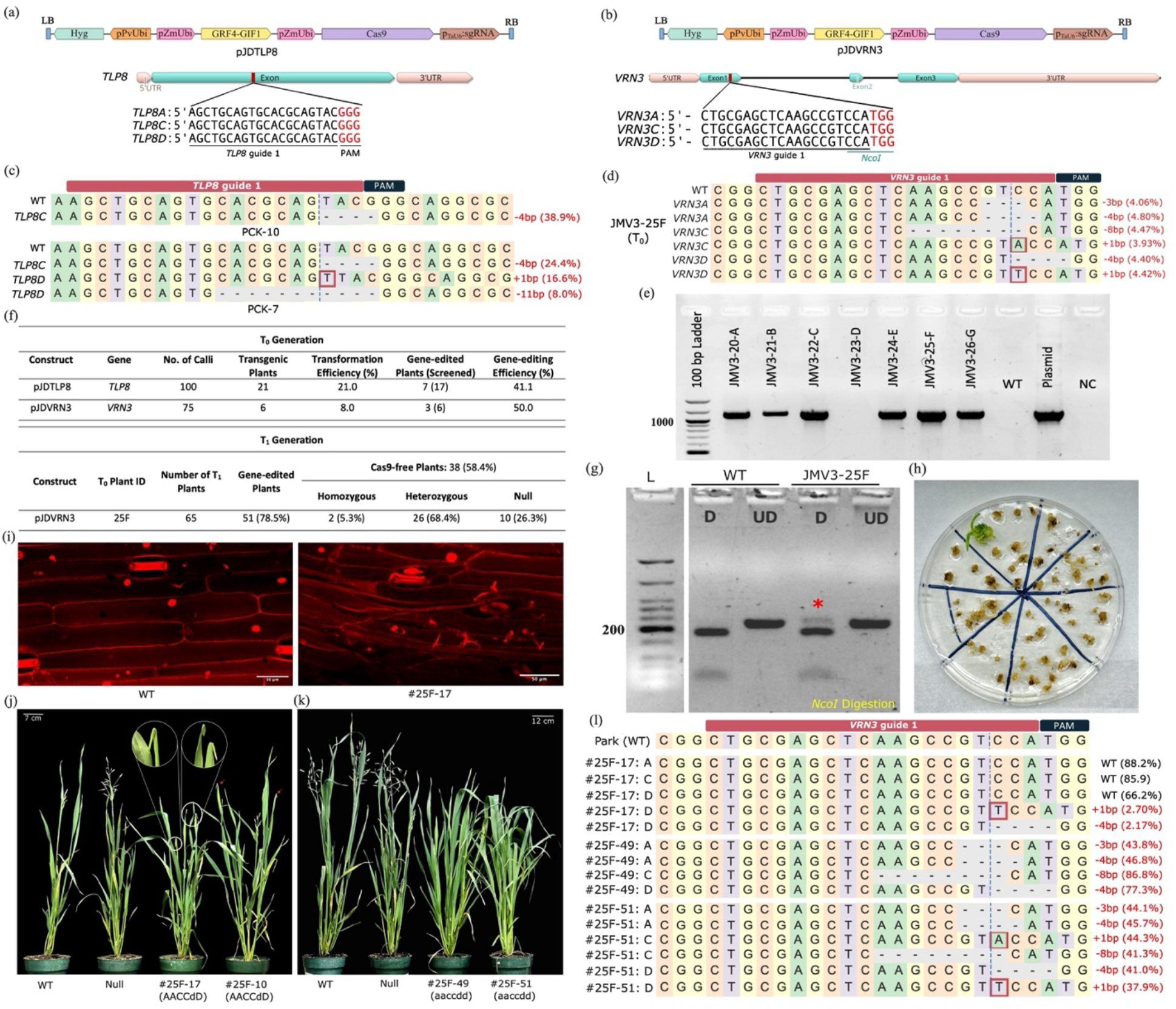
Targeted CRISPR/Cas9-mediated gene-editing in oat. (a) CRISPR construct pJDTLP8 with *TLP8* gRNA from the conserved region. (b) CRISPR construct pJDVRN3 with *VRN3* gRNA from the conserved region. PAM is depicted in red. (c) Targeted mutagenesis in the *TLP8* guide region. Deletions are depicted with a dashed line and insertion with a red box leading to a frameshift mutation. Mutation types are indicated in red on the right. (d) Targeted mutagenesis in the *VRN3* guide region. Deletions are shown with a dashed line and insertion with a red box leading to a frameshift mutation. Mutation types are indicated in red on the right. (e) PCR screening with *hpt* gene primers in pJDVRN3 transformed lines. WT: non-transformed control plant; NC: negative control without DNA. (f) Summary of transformation frequency, gene-editing efficiency and transgenerational inheritance. (g) Cleaved amplified polymorphic sequence (CAPS) assay with *NcoI* for screening of gene-edited pJDVRN3 T_0_ plants. The red star indicates an undigested amplicon emerged due to gene editing. (h) Hygromycin selection (20mg/L) of transgenic calli. (i) Maximum intensity projected para-dermal view of WT and T_1_ line #25F-17 flag leaf segments stained with propidium iodide. Bars = 50 μm. (j) Phenotyping of *VRN3* single copy knockout T_1_ mutants. Bend flag leaves in the heterozygous line #25F-17 are magnified on the top, while for #25F-10, they are indicated with red arrows. Scale bar = 7 cm. (k) Phenotyping of *VRN3* triple knockout T_1_ mutants. Scale bar = 12 cm. (l) Zygosity confirmation of *VRN3* gene-edited T_1_ plants via NGS. Deletions are shown with a dashed line and insertions with a red box. Base pair alterations are indicated in red on the right.

In another experiment, 75 calli were bombarded with pJDVRN3 targeting the oat *VRN3*, yielding six transgenic plants confirmed through *hygromycin (hpt)* gene PCR with a mean transformation efficiency of 8% (Figure 1e,f; Table S1). All the plants were successfully regenerated and transferred to soil in the growth chamber. The target region was PCR amplified from the T_0_ transgenic lines using the *VRN3* primers (Table S1) and sequenced by NGS. The sequencing results reported gene-edited plants with small deletions and insertions in the *VRN3* gene (Figure 1d). Intriguingly, the 4bp deletion in *VRN3D* altered the *NcoI* restriction site that facilitated the screening of knockout mutants through cleaved amplified polymorphic sequence (CAPS) assay. The CAPS genotyping identified the gene-edited lines with 4bp deletion depicting undigested mutated PCR amplicon, while the WT control was completely digested (Figure 1g). The undigested amplicon was gel extracted and 4bp deletion was confirmed by Sanger sequencing (Figure S4). In total, three gene-edited plants were obtained in the T_0_ generation, with mutations in all three *VRN3* copies reporting a high gene editing efficiency of 50 % (Figures 1d,f).

Since we were anticipating a perceptible phenotype in the *VRN3* lines, they were advanced to the next generations. To test the heritability and transgene segregation, 65 T_1_ plants of the JMV3-25F T_0_ line were screened at the guide region using the CAPS assay, followed by NGS and Sanger sequencing. DNA was amplified from 65 T_1_ lines using VRN3D-specific gene primers followed by restriction digestion with *NcoI* as the restriction site has been altered in *VRN3D*-edited plants (Table S1). Of the 65 T_1_ plants screened using CAPS assay, 51 78.5%) contained gene edits for *VRN3D* (Figure 1f and Figure S2). Further analysis revealed 38 Cas9-free plants, out of which two were homozygous, while 26 plants had heterozygous mutations (Figure 1f and Figure S3). The plants were phenotyped in a controlled growth chamber under long-day conditions (16h light/8h dark). Two distinct and intriguing phenotypes were observed in the T_1_ progeny, and their genotypes were extrapolated through NGS using homoeologous gene-specific primers. One set of plants (#25-17, #25F-10, and #25F-39) displayed impaired flag leaf development (sharp bend near the flag leaf tip) at the booting stage (Z41) as compared to the WT and null plants that showed healthy upright flag leaves (Figure 1j) (Zadoks et al., 1974). Such plants carried a heterozygous mutation in the *VRN3D*, while the *VRN3A* and *VRN3C* remained unedited (AACCdD) (Figure 1l, Figure S2, S4). Transgenerational inheritance of the bent leaf trait was consistently observed in the T_2_ generation of the #25F-17 plant. This phenotype was further validated in other *VRN3D* mutants generated through a *VRN3D*-specific guide (Figure S5). The bend site was further investigated under a confocal microscope, revealing changes in the tissue arrangement and morphology of epidermal cells in the mutant compared to the wild type (Figure 1i). Since *VRN* genes belong to the PEBP family, we speculate that the PEBP protein may be involved in controlling plant epidermal cell patterning and differentiation, as observed in other organisms (Trakul et al., 2005). On the contrary, several plants exhibited only a vegetative growth phase and were genotypically characterized as triple knockouts with biallelic or homozygous mutations in all *VRN3* copies (aaccdd) (Figure 1k,l). However, it remains to be seen whether the conserved role of the *SPL*/miR156 module in inflorescence development and reproductive phase transition in oat has been affected (Mehtab-Singh et al., 2024).

In summary, we report the first successful CRISPR-Cas9-based gene editing in oat in three different genes — *AsTLP8, AsVRN3* and *AsVRN3D* with high gene-editing efficiencies. The gene-edited plants for all the genes carried deletions and/or one base insertion. Analysis of *VRN3* mutant T_1_ and T_2_ plants revealed bent leaves in single-copy knockouts (AACCdD), while an extended vegetative growth phase was seen in the T_1_ triple-knockout mutants (aaccdd), accentuating the important role of *VRN3* in oat development. We are confident that this highly efficient oat gene editing system will pave the way for a deeper molecular understanding of this healthy cereal, deciphering oat’s functional genomics, and creating genetic diversity at the cold spots of recombination in oat.

## Author contributions

JS and M-S designed the experiments. M-S and CK designed the gRNA and constructs. M-S, CK and RK performed the oat transformations and tissue culture. M-S and CK conducted the molecular and phenotypic screening. M-S made the figures and wrote the manuscript, and all authors revised it.

## Acknowledgments

This study was financially supported by the Prairie Oat Growers Association (POGA) through Agriculture Funding Consortium. We also acknowledge the support of the NSERC-CREATE program on Genome Editing for Food Security and Environmental Sustainability (GEFSES). We sincerely thank McGill University ECP3-Multi-Scale Imaging Facility, Sainte-Anne-de-Bellevue, Canada and especially Diksha Bhola (McGill University, Montreal) for her assistance with sample preparation and confocal microscopy imaging.

## Supporting Information

**Table S1** List of primers used in the study.

**Methods S1** Material and Methods

**Figure S1** Confirmation of transgenic lines in pJDTLP8 experiment.

**Figure S2** CAPS assay on T_1_ transgenic lines from JMV3-25F.

**Figure S3** Screening of Cas9-free plants in the JMV3-25F T_1_ generation.

**Figure S4** Sanger sequencing of *VRN3* mutant lines.

**Figure S5** Targeted gene editing in the *VRN3D* gene.

