## Supporting Information for "A highly efficient CRISPR-Cas9-based gene editing system in oat (*Avena sativa*)"

Brief Communication

### Methods S1

#### Plant material and callus production

Mature seeds of the oat spring variety *Park* were sterilized in 20% bleach for 20 minutes, followed by sterile water washings. The seeds were put on MS-based (Murashige and Skoog, 1962) regeneration media for germination under low light conditions (16h light, 10-40 µE) at 24 °C. Upon germination, they were placed on modified callus induction media (C') after trimming the grown shoot/roots. After a couple of weeks, superior quality green callus with embryogenic patterns was selected for particle gun bombardment.

#### Guide RNA and binary construct design

The oat genome of cultivar *Park* was downloaded from GrainGenes (https://wheat.pw.usda.gov/GG3/) and uploaded as a custom database to the Geneious Prime software for gRNA design and off-target analysis. The gene sequences of oat *TLP8 (A/C/D)* and *VRN3 (A/C/D)* copies were extracted and aligned. The guides were selected based on a high specificity score (Hsu et al., 2013) and a high activity score (Doench et al., 2014). The guides were separately cloned in the JD633 backbone with the GRF-GIF system described by (Debernardi et al., 2020), yielding three different constructs (pJDTLP8, pJDVRN3, and pJDVRN3D). The oligo pair was specifically designed for the target sequence in a way that created a four-base overhang at both ends (5' –acctg; 3' caaa). The T4 polynucleotide kinase was used to phosphorylate the oligos and hybridized to create a spacer dimer. It was cloned at the *AarI* sites of the binary vector to generate the final CRISPR constructs. *E.coli* DH5α cells were used to transform the final constructs and then grown on selective media (Lysogeny broth (LB) supplemented with specific antibiotic. The QIAGEN Plasmid Midiprep Kit was used to perform plasmid isolation after picking the positive colonies. The final constructs were sequenced to confirm the integration of the guide.

**Genetic transformation and tissue culture**

Before bombardment, a 3 cm diameter circle of callus is initially put on osmotic media for three hours (C' supplemented with 0.2 M mannitol and 0.2 M sorbitol). 6µl of suspended gold stock solution was utilized per plate (60mg of 0.6 µm gold particles in 1ml of 100% ethanol). The supernatant was discarded after centrifuging the suspension for 1 minute at 13,000 rpm. The pellet was again centrifuged after adding 200-300 µl of filter sterile water ($FSH_2O$) and the supernatant was discarded. Again, the pellet was obtained and 6 µg of plasmid DNA was added. Following this, 250 µl of $FSH_2O$ from the volume added in DNA, 250 µl of calcium chloride ($CaCl_2$) and 50 µl of spermidine is added. This mixture was briefly vortexed and incubated on ice for 30 minutes. Then the mixture was again centrifuged at 13,000 rpm for 1-2 minutes and 200 µl of ethanol was added after removing the supernatant. Finally, 36 µl of 95% (v/v) ethanol was added to the pellet to make the plasmids ready for bombardment. The microprojectile bombardment was accomplished using the BioRad PDS-1000/He system (Hercules, CA) at 1100 psi. The next day the bombarded calli were transferred to the modified C' media for a week under low light at 27°C. Three rounds of selections were performed by placing the bombarded callus on modified C' media supplemented with 20 mg/L of hygromycin B (Phytotech) at 27 °C under 16h light/ 8hr dark conditions. The healthy calli were placed on the same media and the process was continued for a few weeks. The selected calli were allowed to regenerate, producing shoots, followed by rooting in the rooting media. Later, they were transferred to pots in the greenhouse after a week of hardening. The plants were grown to maturity in the greenhouse under 16h daylight at 1000 µE and 8h darkness.

**Components of regeneration, rooting and C' Media**

| **Components** | Regeneration | Rooting | C' |
|---|---|---|---|
| MS salts (g/L) | 4.4 | 4.4 | 4.4 |
| Maltose (g/L) | -- | -- | 30 |
| Sucrose (g/L) | 30 | 30 | -- |
| Casein hydrolysate (g/L) | -- | -- | 1 |
| Proline (g/L) | -- | -- | 0.69 |
| Myo-inositol (g/L) | -- | -- | 0.25 |
| Thiamine HCl (mg/L) | 1 | 1 | 1 |
| Pyridoxine HCl (mg/L) | 0.5 | 0.5 | -- |

| | | | |
|---|---|---|---|
| Nicotinic acid (mg/L) | 0.5 | 0.5 | -- |
| CuSO4 | 0.16 mg/L | 0.16 mg/L | 5 µM |
| 2,4-D (mg/L) | | | 2 |
| BAP (mg/L) | 1 | -- | 0.5 |

**Genotyping and mutant identification**

The genomic DNA was extracted from the $T_0$ and $T_1$ plants at (3-4 leaf stage) using the urea method (Chen and Dellaporta, 1994). The GoTaq® Green Master (Promega Corporation, Canada) was used to amplify the incorporated construct by *hygromycin* gene primers (1 µM). The PCR was carried out using the profile: 95 °C for 2 min, followed by 30 cycles of 95 °C for 30 s, 60 °C for 1 minute, 72 °C for 60 s and a final extension at 72 °C for 5 minutes. Gel electrophoresis was carried out to analyze the PCR results. Whereas, the gene sequences flanking guide region was amplified using Q5 High-Fidelity 2X Master Mix (New England Biolabs) following the manufacturer's protocol at an annealing temperature of 60 °C. The transgenic lines and WT control PCR product were subjected to next-generation sequencing, CAPS assay, and Sanger Sequencing for mutant identification. Illumina paired-end read amplicon sequencing and Oxford Nanopore Sequencing were performed by CCIB DNA Core (Cambridge, MA, USA) and Plasmidsaurus (Eugene, OR, USA). NGS raw reads were analyzed using the default parameters in CRISPRESSO2 (Clement et al., 2019) and Geneious Prime software. For the CAPS Assay, the PCR products were subjected to overnight *NcoI* restriction digestion at 37°C, followed by migration at 2% agarose gel. Sanger sequencing was also performed to confirm the gene edits. Sanger trace data was also inferred using the CRISPR Synthego ICE v2 tool (Conant et al., 2022).

**Confocal laser scanning microscopy**

The confocal microscopy imaging was conducted at the McGill University ECP3- Multi-Scale Imaging Facility, Sainte-Anne-de-Bellevue, Quebec, Canada. Small segments from the bend region of the flag leaves at the Z-41 stage were cut. Corresponding regions from the wildtype flag leaf at the same stage were also excised. Both the segments were stained with 1.5 mM propidium iodide for 1.5 hours on rotation. The stained samples were observed under Zeiss LSM710 – Airyscan microscope (Zeiss, Germany) equipped with 20x Plan Apochromat, NA0.8 objective. The samples were excited with a 543 nm (HeNe, 1mW) laser and the images were processed at similar settings using Fiji software (Schindelin et al., 2012).

**Table S1. List of primers used in the study.**

| Primer Name | Description | Sequence |
|---|---|---|
| DeepSeq-AsVRN3A-F | PCR primer to amplify oat *VERNALIZATION 3-A (AsVRN3A)* for genotyping | CGTCAACCCCTAGCATACGAT |
| DeepSeq-AsVRN3C-F | PCR primer to amplify oat *VERNALIZATION 3-C (AsVRN3C)* for genotyping | AGCTGCTAGCTCGCTCCG |
| DeepSeq-AsVRN3D-F | PCR primer to amplify oat *VERNALIZATION 3-D (AsVRN3D)* for genotyping | TCGCTGTTCCAGGCAGCATA |
| DeepSeq-AsVRN3-F | PCR primer to amplify oat *VERNALIZATION 3 (AsVRN3)* for genotyping | GATATGGCCGGGAGGGACAG |
| DeepSeq-AsVRN3-R | PCR primer to amplify oat *VERNALIZATION 3 (AsVRN3)* for genotyping | CGAGTGTGTAGAAGGTCCTCATC |
| DeepSeq-TLP8-F | PCR primer to amplify oat *TLP8* for genotyping | GAAGCTCGACCCGGGGCA |
| DeepSeq-TLP8-R | PCR primer to amplify oat *TLP8* for genotyping | GGGCACGTTGAAGCCGTCGA |
| pRGEB32_7045F | PCR primer to check the *hygromycin (hpt)* presence [1] | TGCTCAACACATGAGCGAAACC |
| pRGEB32_8155R | PCR primer to check the *hygromycin (hpt)* presence [1] | TGAACTCACCGCGACGTCTGTC |
| JD-pZmUbi-F | PCR primer to check the *Cas9* presence [1] | CGATGCTCACCCTGTTGTTTGG |
| JD-Cas9-R | PCR primer to check the *Cas9* presence [1] | GCCAGAGGCGTTGATCGGGTT |
| VRN3 guide 1-F | Forward oligo for *AsVRN3* sgRNA | ACTTGCTGCGAGCTCAAGCCGTCCA |
| VRN3 guide 1-R | Reverse oligo for *AsVRN3* sgRNA | AAACTGGACGGCTTGAGCTCGCAGC |
| VRN3D guide 1-F | Forward oligo for *AsVRN3-D* sgRNA | ACTTGAAGCCGTCCATGGTTGAGGT |
| VRN3D guide 1-R | Reverse oligo for *AsVRN3-D* sgRNA | AAACTACCTCAACCATGGACGGCTTC |
| TLP8 guide 1-F | Forward oligo for *AsTLP8* sgRNA | ACTTGAGCTGCAGTGCACGCAGTAC |
| TLP8 guide 1-R | Forward oligo for *AsTLP8* sgRNA | AAACGTACTGCGTGCACTGCAGCTC |

[1] Biswal, A.K., Hernandez, L.R.B., Castillo, A.I.R., Debernardi, J.M., Dhugga, K.S., 2023. An efficient transformation method for genome editing of elite bread wheat cultivars. Front Plant Sci 14.

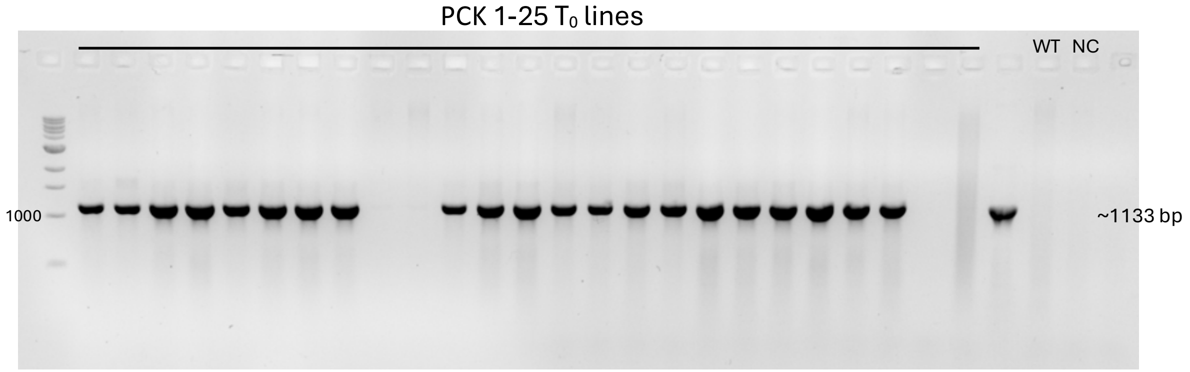


**Figure S1.** Confirmation of $T_0$ transgenic lines transformed with the pJDTLP8 construct using *hpt* gene primers. WT: non-transgenic Park; NC: no template control

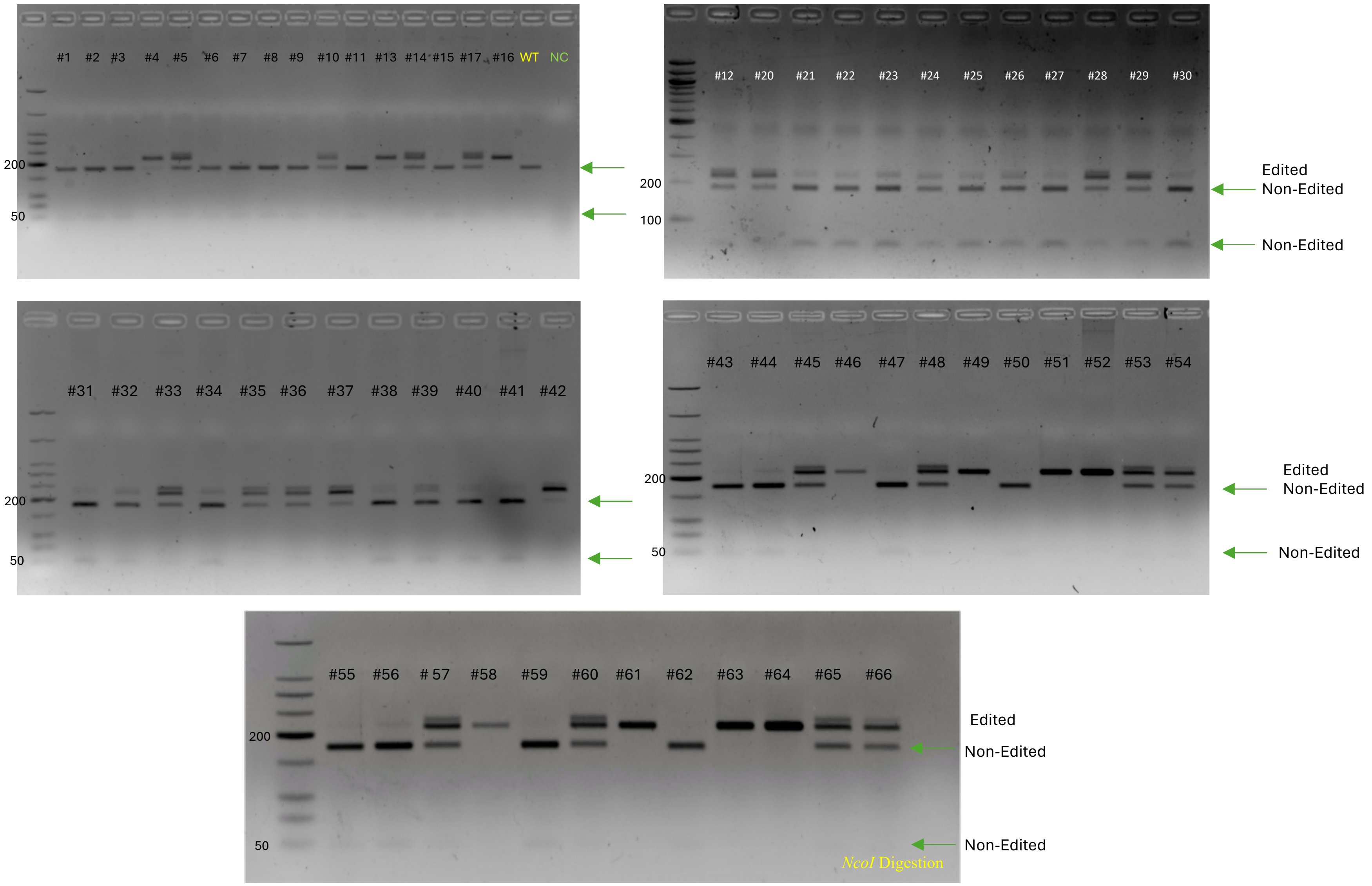


**Figure S2.** CAPS assay on $T_1$ transgenic plants from JMV3-25F. Editing disrupts an *NcoI* restriction site in the target region, resulting in an undigested band (red arrow), while the non-edited sequences are digested. WT- non-transgenic Park; NC: no template control

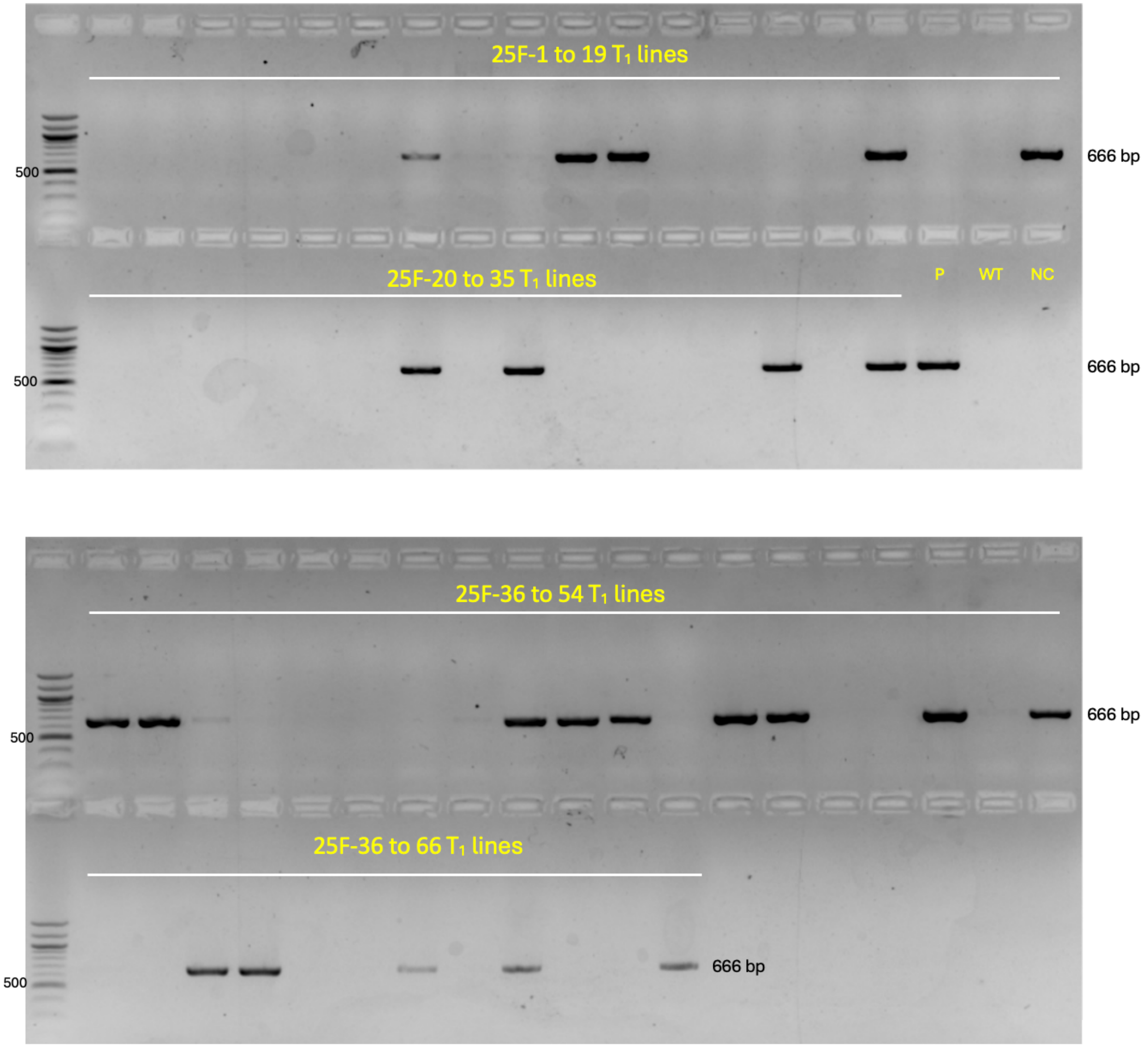


**Figure S3.** Screening of Cas9-free plants in the JMV3-25F $T_1$ generation. Faint bands were also considered Cas9 positive. P: Plasmid; WT: non-transgenic Park; NC: no template control

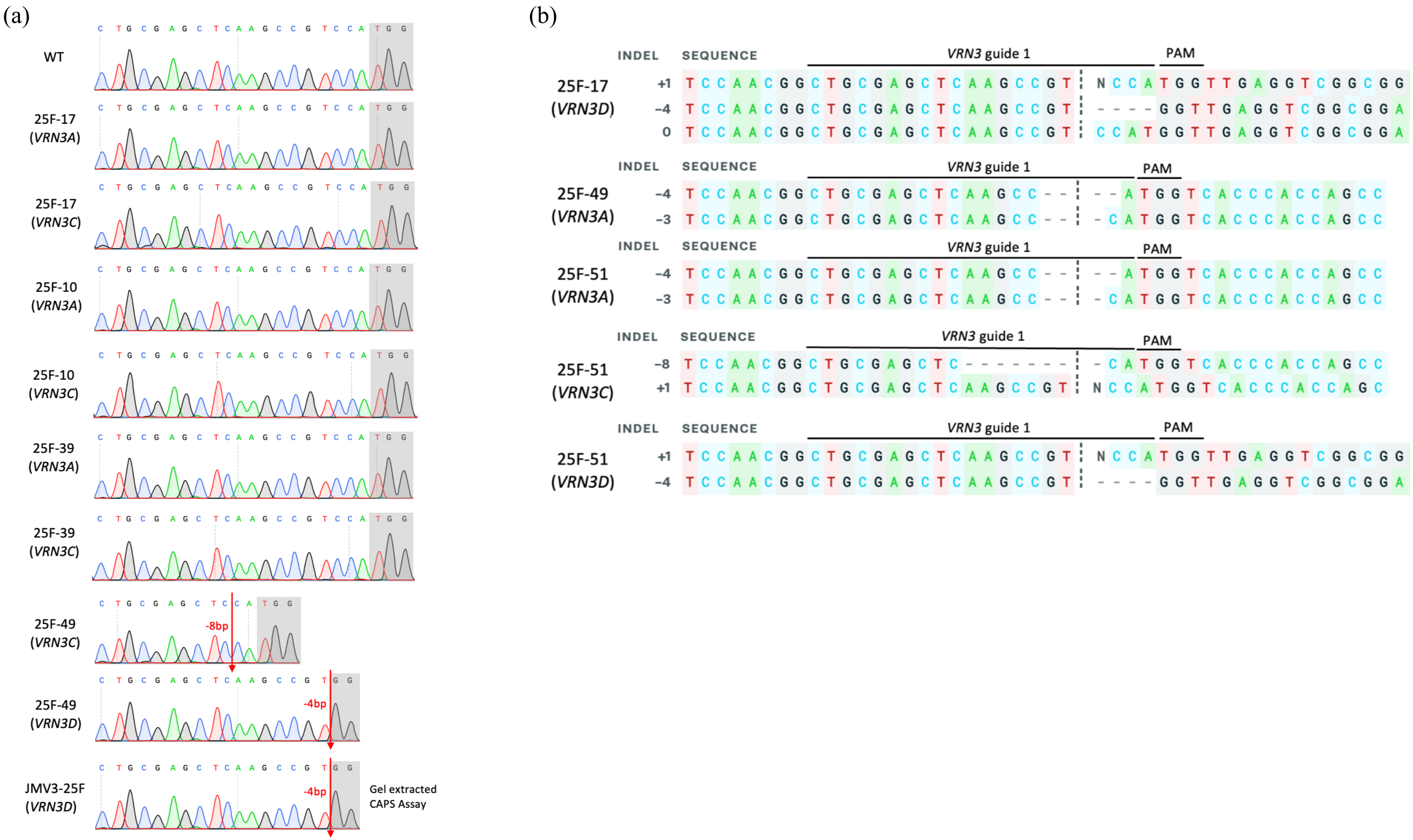


**Figure S4.** Sanger sequencing of *VRN3* mutant lines with homoeologous specific primers. (a) WT and mutant chromatograms are shown at the guide region. PAM is highlighted in grey, and the deletions are depicted with a red arrow. (b) Sequence analysis of *VRN3* target region in $T_1$ biallelic and heterozygous lines using the Synthego ICE v2 tool.

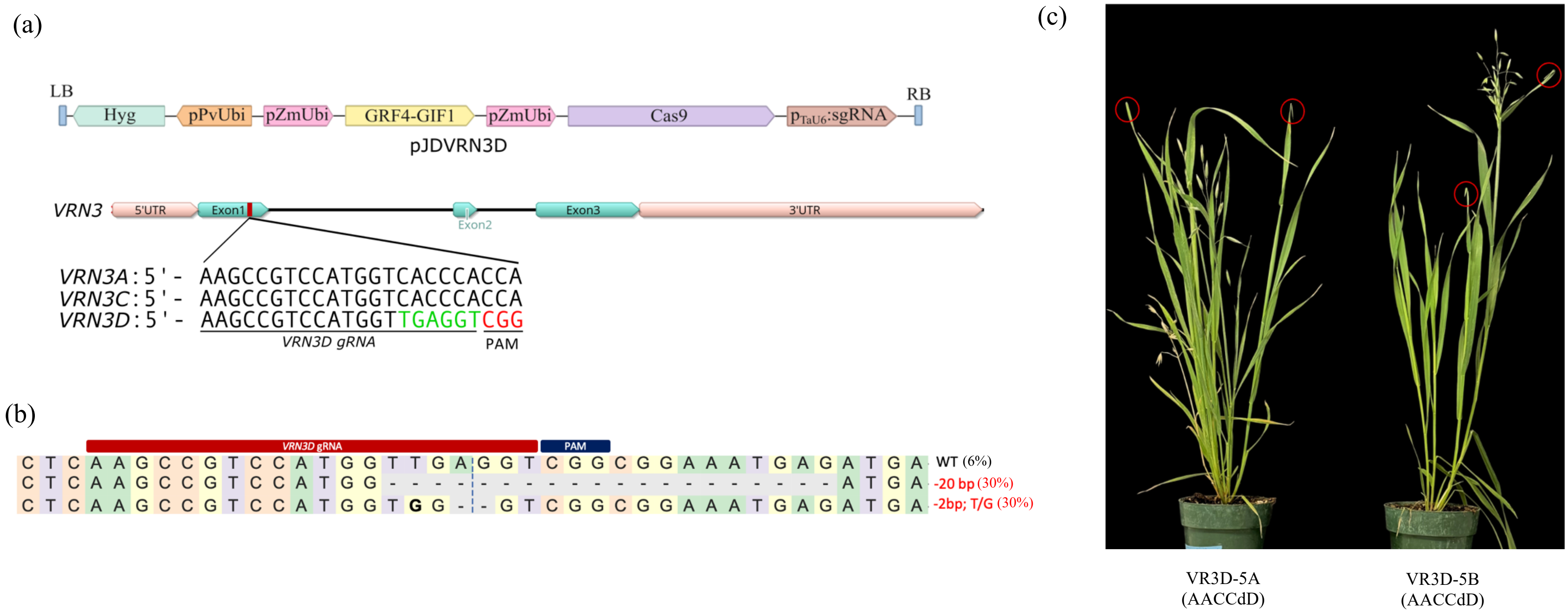


**Figure S5.** Targeted gene editing in the *VRN3D* gene. (a) CRISPR construct pJDVRN3D with *VRN3D*-specific gRNA. (b) Targeted mutagenesis in the *VRN3D* guide region. Deletions are shown with a dashed line. Mutation types are indicated in red on the right. (c) Confirmation of bent flag leaf phenotype in the $T_0$ lines derived from pJDVRN3D, specifically targeting the *VRN3D* copy. Bent flag leaves are circled in red.
